# Phagocytosis by stroma confounds co-culture studies

**DOI:** 10.1101/2020.12.11.417923

**Authors:** Sophie A. Herbst, Marta Stolarczyk, Tina Becirovic, Yi Liu, Carolin Kolb, Marco Herling, Carsten Müller-Tidow, Sascha Dietrich

## Abstract

Signals provided by the microenvironment can modify and circumvent pathway activities that are therapeutically targeted by drugs. Bone marrow stromal cell co-culture models are frequently used to study the influence of the bone marrow niche on ex-vivo drug response. Here we show that mesenchymal stromal cells from selected donors and NKTert, a stromal cell line which is commonly used for co-culture studies with primary leukemia cells, extensively phagocytose apoptotic cells. This could lead to misinterpretation of the results, especially if the viability readout of the target cells in such co-culture models is based on the relative proportions of dead and alive cells. Future co-culture studies which aim to investigate the impact of bone marrow stromal cells on drug response should take into account that stromal cells have the capacity to phagocytose apoptotic cells.

## Introduction

Signals provided by the microenvironment protect leukemia cells from spontaneous apoptosis and modify pathway activities which are therapeutically targeted by drugs (Choi et al., 2016; Ten Hacken and Burger, 2016). *In-vitro* co-culture models of malignant cells and various bone marrow stromal cell types have been frequently used to understand tumor - microenvironment interactions and how these interactions interfere with the response to cancer drugs. Most commonly, co-cultures of leukemia cells and the human bone marrow stromal cell lines NKTert and HS-5, or primary mesenchymal stromal cells (MSCs) are used as models of the human bone marrow niche. Several reports suggest that NKTert and other stromal cells protect leukemia cells from spontaneous and drug induced apoptosis (Balakrishnan et al., 2015; Cheng et al., 2014; Fiorcari et al., 2013; Kurtova et al., 2009; Zhang et al., 2012). Here we demonstrate that NKTert and primary MSCs from selected donors massively phagocytose apoptotic cells *in-vitro*. The clearance of dead cells increases the relative proportion of alive cells in these co-cultures, which represents a major source of bias if the viability readout is based on relative proportions of alive and dead cells. In contrast, other co-cultures (with some MSCs or HS-5) did not show this behaviour, suggesting them as suitable feeder cells for co-culture drug response studies.

## Results and Discussion

Cancer cells from four CLL patients were cultured either alone (mono-cultures) or in co-culture with the NKTert stromal cell line. The mono- and co-cultures were treated with solvent control (DMSO), venetoclax or fludarabine. After 72 hours, the cultures were stained with the nuclear dye SiR-DNA, the viability dye calcein and the dead cell marker propidium iodide (PI). Based on these stainings we quantified dead and alive CLL cells by flow cytometry and high content confocal microscopy. First we calculated relative proportions of dead and alive cells. In accordance with the literature (Balakrishnan et al., 2015; Cheng et al., 2014; Fiorcari et al., 2013; Kurtova et al., 2009; Zhang et al., 2012) we could confirm higher proportions of alive CLL cells in co-cultures with NKTert stromal cells, than in suspension mono-cultures of CLL cells only. This was true for all untreated and drug treated conditions and could be interpreted as stromal cell mediated protection from spontaneous and drug-induced apoptosis. We further compared absolute CLL cell counts as determined by microscopy and flow cytometry. Surprisingly, we observed much lower absolute CLL cell counts in NKTert co-cultures than in CLL mono-cultures (Fig. 1b). This decrease was due to the disappearance of dead cells (Fig. 1c, left). Absolute counts of alive cells in the cultures did not differ between mono- and co-cultures with NKTert (Fig. 1c, right).

**Fig. 1:**
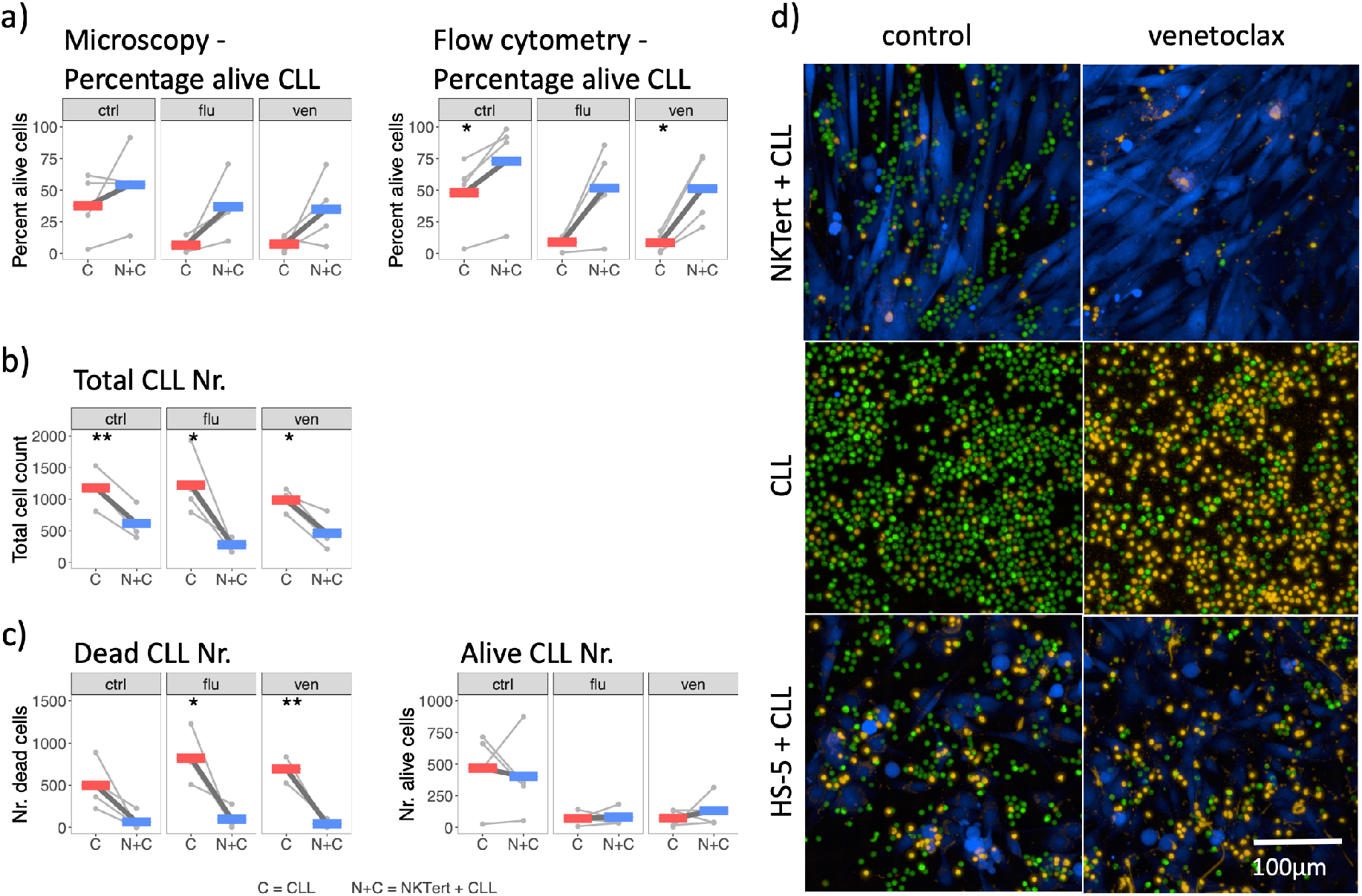
Dead leukemia cells disappear from co-cultures with NKTert bone marrow stromal cells. **a,** The percentage of alive CLL cells was higher in co-cultures of CLL and NKTert (N+C) than in CLL mono-cultures (C). This was true for cultures treated with solvent (ctrl), fludarabine (flu) or venetoclax (ven). The readout was either performed with microscopy or flow cytometry; paired t-test, * = p < 0.05; thin lines = samples from four patients; thick, coloured bars = mean. **b,** Total cell counts decreased in solvent control treated (ctrl), fludarabine treated (flu) and venetoclax treated (ven) co-cultures of CLL and NKTert (N+C) in comparison to mono-cultures of CLL (C). Paired t-test, ** = p < 0.01, * = p < 0.05; thin lines = samples from four patients; thick, coloured bars = mean, assessed by microscopy. **c,** Dead CLL cells disappeared in co-cultures with NKTert. The total number of alive cells was comparable between the culture conditions. Paired t-test, ** = p < 0.01, * = p <0.05; thin lines = samples from four patients; thick, coloured bars = mean, assessed by microscopy. **d,** Confocal microscopy pictures of CLL cells cultured in mono-cultures or in co-culture with NKTert or HS-5 stromal cells. CLL cells disappeared from CLL-NKTert co-cultures treated with venetoclax. Green = CellTracker Green, blue = CellTracker Blue, yellow = propidium iodide.

To better understand this finding, we labeled leukemia cells from four other CLL patients with CellTracker Green and co-cultured them with CellTracker Blue labelled NKTert cells (Fig. 1d). Venetoclax pretreated and apoptotic leukemia cells disappeared already after 16 hours in co-culture with NKTert cells, while apoptotic cells were still present in mono-cultures. We further aimed to understand whether other stromal cells behave in the same way and co-cultured venetoclax pretreated CLL cells with the cell line HS5. Cell counts of CLL cells were comparable between mono-cultures of CLL cells and co-cultures with HS5 stromal cells. These experiments indicated that apoptotic leukemia cells disappeared in NKTert, but not in HS-5 co-cultures, which increased the relative proportion of alive leukemia cells (Fig. 1a) and could falsely be interpreted as protection of CLL cells by NKTert from spontaneous or drug-induced apoptosis.

Next we aimed to identify why apoptotic cells disappeared in NKTert cell line co-cultures. Especially in the venetoclax treated co-cultures many CLL cells seemed to be located inside of NKTert, based on bright field microscopy (Fig. 2a and S1). Confocal microscopy could confirm the localisation of CellTracker Green labeled CLL inside of CellTracker Blue labeled NKTert (Movie M1). Further staining with the lysosomal dye NIR revealed large lysosomal bodies inside NKTert cells (Fig. 2b), which had the size and shape of CLL cells. Lysosomal bodies often occurred in regions also positive for CellTracker Green and PI staining and were surrounded by, but did not include CellTracker Blue staining (Fig. 2c and Movie M1). We quantified the amount of these phagosomes in three independent experiments (Fig. 2d). A high number of phagosomes was present in treated NKTert co-cultures, whereas large lysosomal bodies were absent in CLL mono-cultures, NKTert or HS-5 stroma mono-cultures or co-cultures of CLL and HS-5 cells. We concluded that NKTert cells, but not HS-5, phagocytose apoptotic CLL cells and, thereby, clear those cells from the co-cultures.

**Fig. 2:**
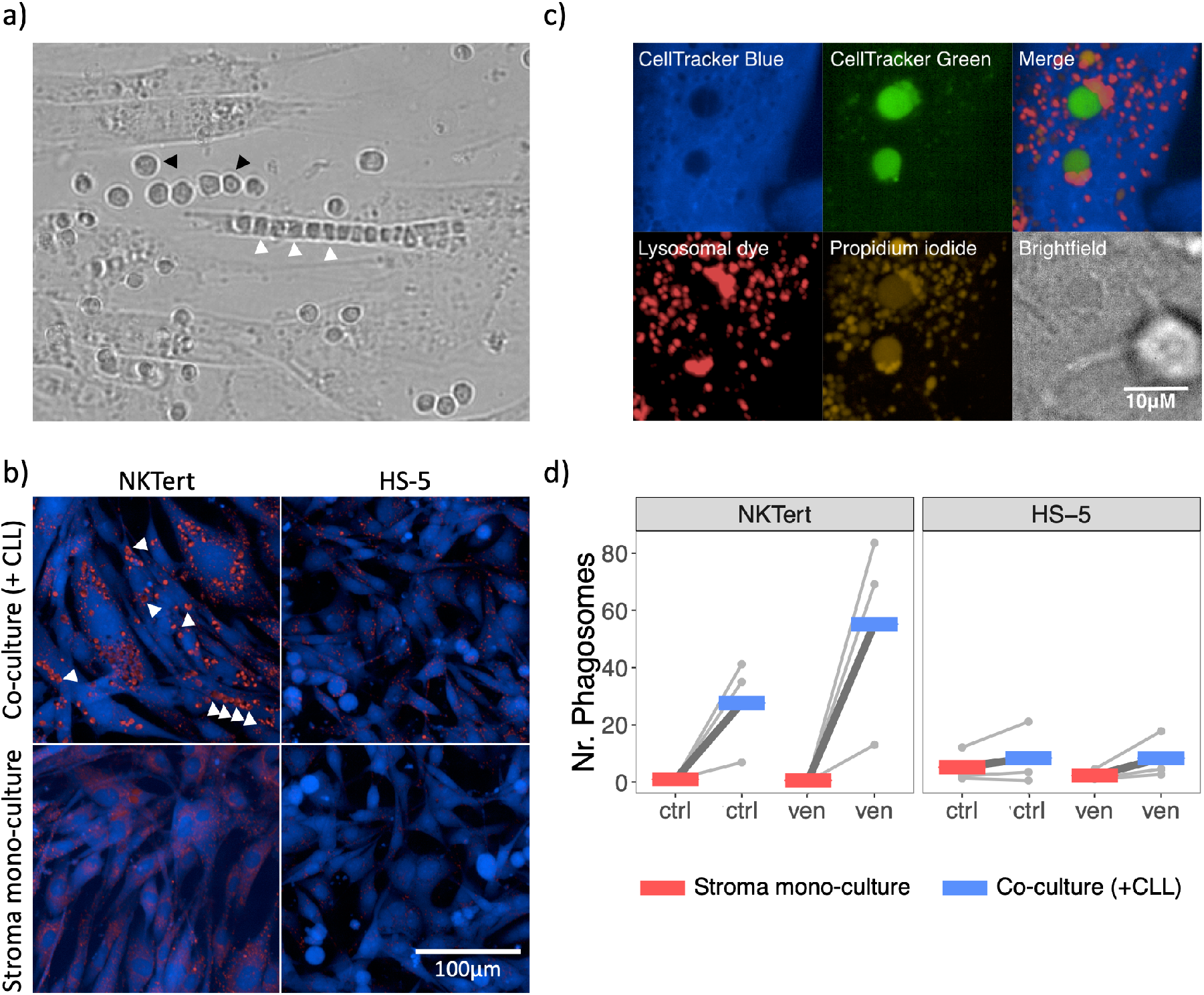
Disappearance of dead leukemia cells is due to extensive phagocytosis by NKTert bone marrow stromal cells. **a,** Brightfield image of venetoclax treated NKTert and CLL co-culture showing a NKTert cell having phagocytosed many CLL cells. Three of the many phagocytose CLL cells are highlighted with white arrows, two non-phagocytosed CLL cells are highlighted with black arrows. **b,** Large lysosomal bodies (phagosomes) appeared in co-cultures of NKTert and CLL treated with venetoclax. Some of the phagosomes are highlighted with white arrows. Blue = CellTracker Blue, red = lysosomal dye NIR. **c,** Magnification of two phagosomes inside NKTert (CellTracker Blue) cells. CLL cells had been previously labelled with CellTracker Green. **d,** Quantification of phagosomes in microscopy pictures of HS-5 or NKTert stroma mono-cultures or co-cultures with CLL cells. ctrl = solvent control, ven = venetoclax; Points and lines = individual patients, coloured bars = mean; samples from three CLL patients.

We wondered whether phagocytosis by NKTert was specific to CLL cells. To this end, the mantle cell lymphoma cell line HBL-2, the acute myeloid leukemia cell line OCI-AML 2, the carcinoma cell line HELA, and the benign epithelial cell line HEK-293T were exposed to 10 µM doxorubicine, to induce apoptosis. Cells were then co-cultured with NKTert cells. NKTert phagocytosed apoptotic cells of all tested cell lines, regardless of their origin (Fig. 3a). In conclusion, phagocytic activity by NKTert cells is not limited to CLL cells, but is likely cell type independent.

**Fig. 3:**
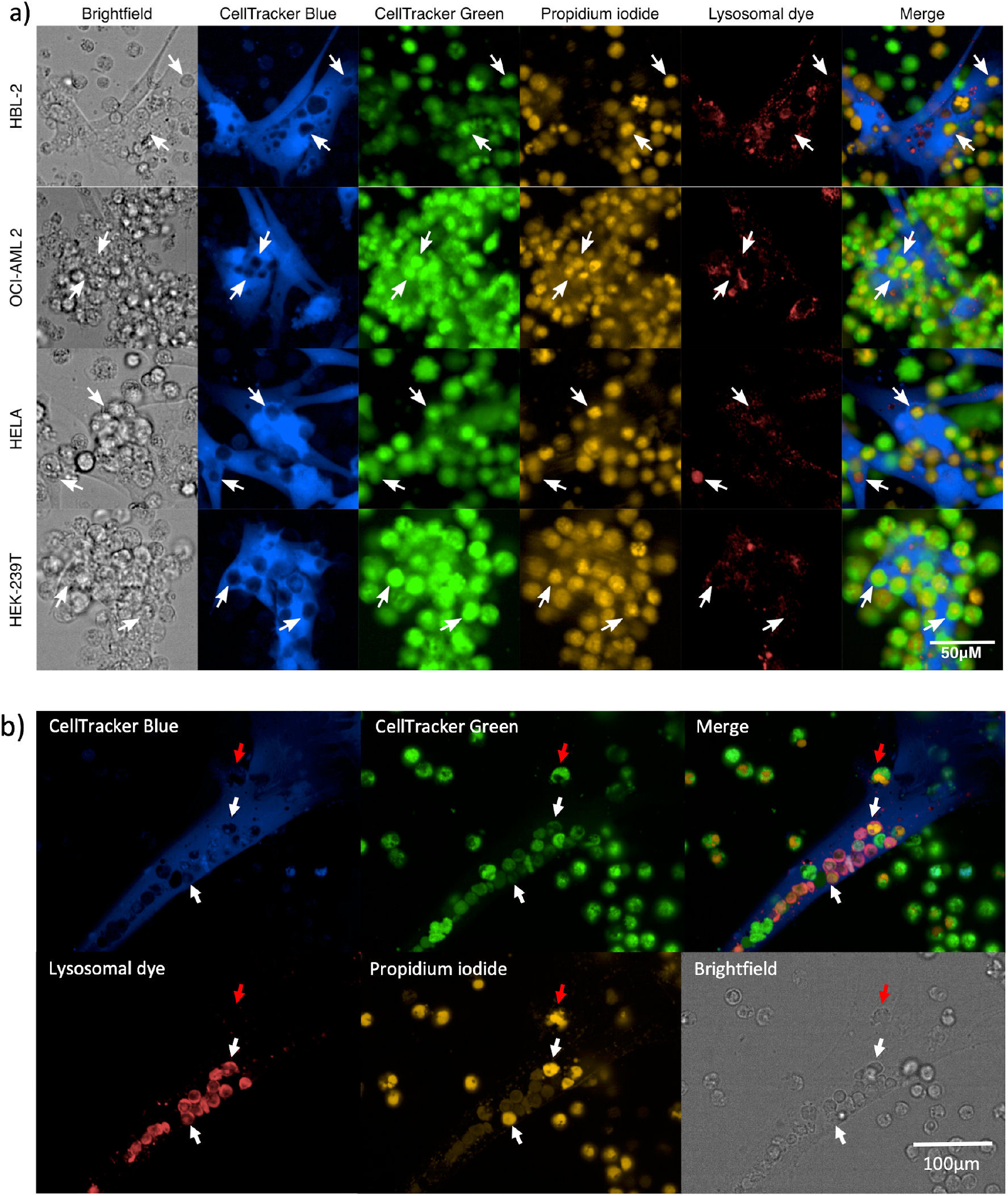
Phagocytosis is not restricted to NKTert or dead leukemia cells. **a,** NKTert also phagocytosed apoptotic cells of non-haematopoietic origin. CellTracker Green labelled HBL-2, OCI-AML 2, HELA or HEK-293T cells (green), treated with doxorubicine to induce apoptosis and co-cultured with NKTert. Blank areas in staining of CellTracker Blue labelled NKTert (blue) were observed (examples indicated by arrows). These blank areas overlapped with CellTracker Green (green) and propidium iodide (yellow) stained HBL-2, OCI-AML 2, HELA or HEK-293T cells. Some of these phagosomes were acidic, indicated by staining with lysosomal dye NIR (red). **b,** CellTracker Blue labelled mesenchymal stromal cell (MSC; blue) phagocytosing dead CellTracker Green labelled CLL cells (green). White arrows indicate examples for phagosomes containing dead CLL cells. Red arrow indicates dead CLL cell in the process of being phagocytosed by NKTert. See also movie M2 for 3D view.

To further evaluate whether the ability to phagocytose apoptotic cells is restricted to the cell line NKTert or whether it also occurs in primary MSCs, which are a more physiological model of the bone marrow niche than cell lines, we cultured venetoclax treated and CellTracker Green labelled primary CLL cells with CellTracker Blue labelled primary MSCs from four different healthy donors. Phagocytosed CLL cells were observed in MSCs from two out of four donors (Fig. S2). Phagocytosis was especially high in MSC1 culture (Fig. 3b, Movie M2), while MSC2 only exhibited a low amount of phagocytosis (Fig. S2). This shows that not only NKTert, but also some, but not all, primary MSCs are able to phagocytose apoptotic cells.

Due to the massive phagocytic activity exhibited by NKTert we assessed whether or not the cells are of macrophagic origin. By performing staining against CD45 followed by flow cytometry we could show that NKTert are negative for this pan-hematopoietic cell marker and, based on this, are not of monocytic lineage or conventional macrophage differentiation (Fig. 4a). This confirmed the findings by Kawano et al. (2003). Therefore, the extensive phagocytosis by NKTert cannot be explained by a macrophagic origin of NKTert.

**Fig. 4:**
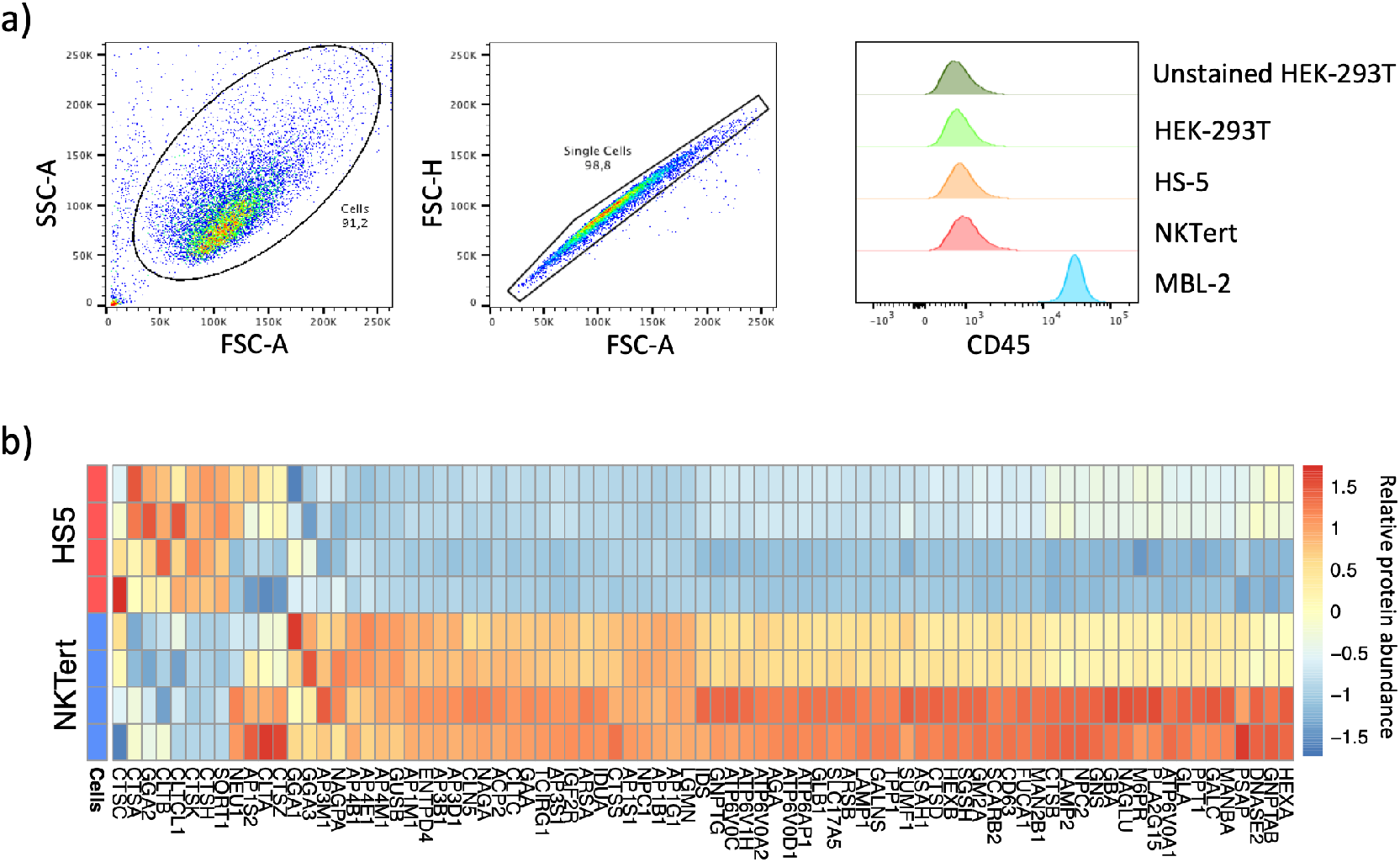
Molecular differences between NKTert and HS-5 bone marrow stromal cell lines. **a,** CD45 abundance was assessed by flow cytometry in NKTert and HS-5 cells and in HEK-293T and MBL-2 control cells. Gating strategy and results are shown. NKTert were CD45 negative, and therefore a hematopoietic and, thus, monocytic or conventional macrophagic origin was excluded. **b,** Relative abundances of proteins in the lysosomal pathway for HS-5 and NKTert stromal cells, assessed by proteomics in four replicates.

Since only one out of the two tested human bone marrow stromal cell lines and two out of the four tested MSCs phagocytosed apoptotic cells, we were interested in molecular differences associated with this phagocytic activity. We performed quantitative proteomics of HS-5 and NKTert cells. Pathway enrichment analysis on protein abundances revealed that the lysosomal pathway was most significantly enriched within differential proteins. Most proteins in this pathway were upregulated in NKTert in comparison to HS-5 (Fig. 4b). This indicates that upregulation of lysosomal proteins contributes to the phagocytic activity of NKTert. Ding et al. (2018) reported that blocking of the lysosomal pathway leads to the presence of more apoptotic CLL cells in NKTert co-cultures. However, the authors interpreted this finding as a disruption of the protection from apoptosis by NKTert, rather than an inhibition of phagocytic activity.

## Conclusion

In conclusion, we found that NKTert and selected primary MSCs massively phagocytose apoptotic cells *in-vitro*. Although it has been reported that MSCs are capable of phagocytosis (Dogusan et al., 2004), the extent and consequences for *in-vitro* assays have not been anticipated. Phagocytosis leads to the removal of dead cells from the co-cultures causing an increased percentage of alive cells, which might be falsely interpreted as a protective effect mediated by stromal cells. This bias might affect flow cytometry assays which rely on the measurement of the relative proportion of alive and dead cells without assessing total cell counts in an *in-vitro* culture model. Together this suggests that results from co-culture experiments with NKTert or selected MSCs could be misinterpreted if not corrected for this potential bias triggered by the phagocytic activity of these cell types.

There is no doubt that selected bone marrow stromal cells and cell lines are able to protect leukemia cells from spontaneous and drug-induced apoptosis (Lagneaux et al., 1998). For some, but not all, stromal cells this effect might be confounded by their phagocytic activity. The cell line HS-5 for example did not phagocytose apoptotic cells and might therefore be a suitable co-culture model system. For future studies, phagocytosis by stromal cells needs to be tested and taken into account, especially as the heterogeneity between cell lines or donors seems to be high. Even though first approaches into this direction have been taken (Baccin et al., 2020; Baryawno et al., 2019), stromal cells are still not sufficiently characterized. Only this will ensure accurate determination of the influence of the microenvironment and, thus, the development of effective treatment strategies.

## Supporting information

Proteomics NKTert and HS-5

## Acknowledgements

SD was supported by a grant of the Hairy Cell Leukemia Foundation, the Heidelberg Research Centre for Molecular Medicine (HRCMM) and an e:med BMBF junior group grant. CMTs work on MSC was supported by the Deutsche Krebshilfe (70112974) and parts of this work were supported by a grant of the Deutsche Krebshilfe (70172788) to MH. The authors gratefully acknowledge the data storage service SDS@hd supported by the Ministry of Science, Research and the Arts Baden-Württemberg (MWK) and the German Research Foundation (DFG) through grant INST 35/1314-1 FUGG and INST 35/1592-1 FUGG. The authors would like to gratefully acknowledge technical support from the EMBL genomics core facility, the EMBL proteomics core facility and the DKFZ light microscopy facility. We especially thank Peer Haberkant, Frank Stein, Vladimir Benes, Nayara Azevedo and Angela Lenze for technical support.

## Competing interests

The authors declare no competing interests.

## Author contributions

SH, MS, SD planned the experiments. YL, CK, MH provided important input for the experimental designs. SH, MS, TB, CK performed the experiments. SH, MS analysed the data. CMT, SD supervised the work. SH, MS, SD wrote the manuscript. SH and MS contributed equally to this work.

## Supplementary Methods

### Patient samples

Written consent was obtained from patients according to the declaration of Helsinki. Leukemia cells were isolated from blood using Ficoll density gradient centrifugation. Cells were viably frozen and kept on liquid nitrogen until use.

### Cell culture

HS-5 cells were a kind gift by Martina Seiffert. NKTert cells were obtained from RIKEN BRC. Cell lines were maintained in RPMI supplemented with 10 % FBS, 1 % Penicillin/Streptomycin and 1 % Glutamine at 37°C and 5 % CO_2_ in a humidified atmosphere. Primary MSCs were maintained in Bulletkit medium (Lonza). Cell lines were tested for mycoplasma before all experiments using a PCR based testing procedure. Authentication of HS-5 and NKTert cells was performed (Multiplexion). HS-5 cells could be successfully authenticated. For NKTert no reference sequence was available in the database, however, the NKTert DNA was identified as unique sequence, did not match to any other cell line and, therefore, cross contamination could be excluded.

### Microscopy dyes used in this study

In the study the following dyes for microscopy were used at the indicated concentration. The indicated wave lengths were used for fluorophore excitation: Hoechst 33342 (4 μg/ml, Invitrogen, 405 nm), CellTracker Blue (10 μM, Invitrogen, 405 nm), Calcein (1 μM, Invitrogen, 488 nm), CellTracker Green (10 μM, Invitrogen, 488 nm), propidium iodide (PI; 5 μg/ml, Sigma-Aldrich, 561 nm), SiR-DNA (1 μM, Spirochrome, 640 nm), lysosomal dye NIR (1 μl/ml, Abcam, 640 nm). These dyes were used in the following combinations: Hoechst, Calcein, PI and lysosomal dye NIR, or Calcein, PI and SiR-DNA or CellTracker Blue, CellTracker Green, PI and lysosomal dye NIR. With the used setup no crosstalk between the channels was seen, with the exception of a slight spillover between calcein and PI and a slight spillover of the signal from the lysosomal dye into the channel for PI.

### Comparing number and percentages of alive and dead cells in mono- and co-cultures using microscopy and flow cytometry

This section describes the experiment shown in Fig. 1a-c. We cultured cancer cells from four different patients with CLL (2×10^5^/well) either alone or in co-culture with NKTert (1×10^4^/well) in 96-well glass bottom microscopy plates (zell-kontakt GmbH). Stromal cells were seeded 24 hours before the addition of leukemia cells. The samples were treated with solvent control (DMSO), 63 nM venetoclax or 10 μM fludarabine. After 72 hours the cultures were stained with the nuclear dye SiR-DNA, the viability dye Calcein and the dead cell marker PI. Images were taken with the confocal LSM710 microscope (Zeiss) equipped with climate control (37°C, 5% CO_2_) using a 20x objective lens. Z-stack images were acquired in triplicate wells, and within one well 4 adjacent fields were imaged.

Image analyses were performed on all acquired images using KNIME software. Viability of the lymphocytes was calculated based on the Calcein and PI signals in the z-stack images that correspond to live and dead cells, respectively. After background subtraction using a ‘rolling ball’ method (radius: 10) and a global threshold was applied (Huang method). Objects were separated using Watershed. Total cell counts were obtained based on SiR-DNA signal (labeling filter: 25-250 pixels). Live/dead cell populations were classified based on the intensity histogram of Calcein and PI signal with a selection of the threshold separately for each cell culture condition: CLL cell culture, and co-culture of CLL cells with NKtert cells or HS5 cells.

Directly after acquisition of the confocal microscopy pictures, lymphocytes previously labeled with SiR-DNA, Calcein and PI were pipetted from each cultural condition to a 96-well round bottom plate (Greiner) for further analysis with an IQue Screener (Intellicyt). Calcein and PI signals were recorded. The following gating strategy was pursued: Exclusion of potentially remaining stromal cells, by setting of a lymphocyte gate and exclusion of doublets. The percentage of alive cells was determined by gating on Calcein positive and PI negative cells.

### Visualisation of disappearance of leukemia cells and presence of phagosomes by CellTracker and lysosomal staining

This section describes the experiment shown in Fig. 1d-e. We labeled leukemia cells from four other patients with CellTracker Green and co-cultured them (2×10^5^ cells/well) with CellTracker Blue labelled NKTert (1×10^4^ cells/well) or HS-5 (2×10^4^ cells/well) stromal cells. CLL cells were pretreated with solvent control or 63 nM venetoclax for 24 hours. Co-culturing was performed for 16 hours. Cultures were additionally stained with lysosomal dye NIR and PI before imaging. The samples were imaged on an Opera Phenix microscope (Perkin Elmer) in confocal mode. For clearer visualisation, the signal from the lysosomal dye is not shown in Fig. 1d, while Cell Tracker Green signal and PI staining are not shown Fig. 1e. The images shown are representative of all experiments.

### Quantification of the amount of phagosomes

This section describes the experiment shown in Fig. 1f. The amount of phagosomes was quantified in three independent experiments comprising in total three different CLL patient samples. NKTert cells were seeded at a density of 2.5×10^3^ cells/well, HS-5 at a density of 5×10^3^ cells/well into wells of a 384 μ-Clear microscopy plate (Greiner) in technical triplicates. After 4 hours CLL cells were added at a density of 5×10^4^cells/well to co-culture wells, while additional medium was added to stroma mono-culture wells. The cultures were treated with 10 nM Venetoclax or left untreated. After 48 hours the cultures were stained with Hoechst 33342, Calcein, PI and lysosomal dye NIR. Three positions per well were imaged on a CellObserver microscope (Zeiss). The number of phagosomes was quantified in all acquired images using a custom script written in R, with the help of the EBImage package. In brief, a global threshold was applied on the signal from the lysosomal channel, followed by a filling of holes with the *fillHull* function. To separate objects in close proximity a watershed operation was performed on the distance map of foreground and background pixels. Only objects ranging between 100 and 1500 pixels in size and a circularity of more than 0.8 were classified as phagosomes.

### Co-cultures of NKTert and apoptotic cells of different origins

This section describes the experiment shown in Fig. 2a. To investigate whether phagocytosis by NKTert was specific for dead CLL cells, the mantle cell lymphoma cell line HBL-2, the acute myeloid leukemia cell line OCI-AML 2, the carcinoma cell line HELA or the benign epithelial cell line HEK-293T were detached if adherent and labelled with CellTracker Green. The cells were treated with solvent control or 10 μM doxorubicine to induce apoptosis. This was carried out in 15 ml Falcon tubes to avoid reattachment of the adherent cells. NKTert cells were seeded into wells of a 384 μ-Clear microscopy plate (Greiner) at a density of 2.5×10^3^ cells/well and labelled with CellTracker Blue according to the manufacturer’s instructions. After incubation for 24 hours, target cells were added to NKTert cells at a density of 2.5×10^4^ cells/well. Co-cultures of NKTert and CLL cells were used as positive control for the occurrence of phagocytosis. The cultures were stained with PI and lysosomal dye NIR after co-culturing for 16 hours. The samples were imaged on an Opera Phenix microscope (Perkin Elmer) in confocal mode. Representative images are shown.

### Flow cytometry of CD45

This section describes the experiment shown in Fig. 2b. HEK-293T, HS-5, and NKTert cells were harvested with Accutase (Innovative Cell Technologies) to avoid cleavage of surface epitopes. MCL-2 cells were resuspended. Cells were stained with CD45 (BD Pharmingen™ FITC Mouse Anti-Human CD45, Clone: HI30; BD Biosciences) and analysed on a LSRII flow cytometer (BD Biosciences).

### Co-cultures of leukemia cells and primary MSCs using Cell Tracker

This section describes the experiment shown in Fig. 2c. Primary CLL cells were labelled with CellTracker Green. Apoptosis was induced by treatment with 63 nM venetoclax. After 24 hours the cells (2×10^5^ cells/well) were added to primary MSCs of four different healthy donors (1×10^3^ cells/well), labelled with CellTracker Blue into 96-well glass bottom microscopy plates (zell-kontakt GmbH). After incubation for 16 hours the cultures were stained with lysosomal dye NIR and PI. The samples were imaged on an Opera Phenix microscope (Perkin Elmer) in confocal mode. Representative images are shown.

### Proteomics of HS-5 and NKTert

This section describes the experiment shown in Fig. 2d. HS-5 or NKTert cells were trypsinized, washed twice with PBS and snap frozen in liquid nitrogen. Analysis of the whole proteome using mass spectrometry was performed by the EMBL proteomics core facility. Differentially abundant proteins were analyzed using limma (Ritchie et al., 2015). Gene set enrichment analysis for the KEGG pathways (Kanehisa and Goto, 2000) was performed using GSEA (Subramanian et al., 2005). A heatmap of the protein abundance of proteins in the lysosomal pathway was visualised using R.

## Supplementary Figures

**Fig. S1:**
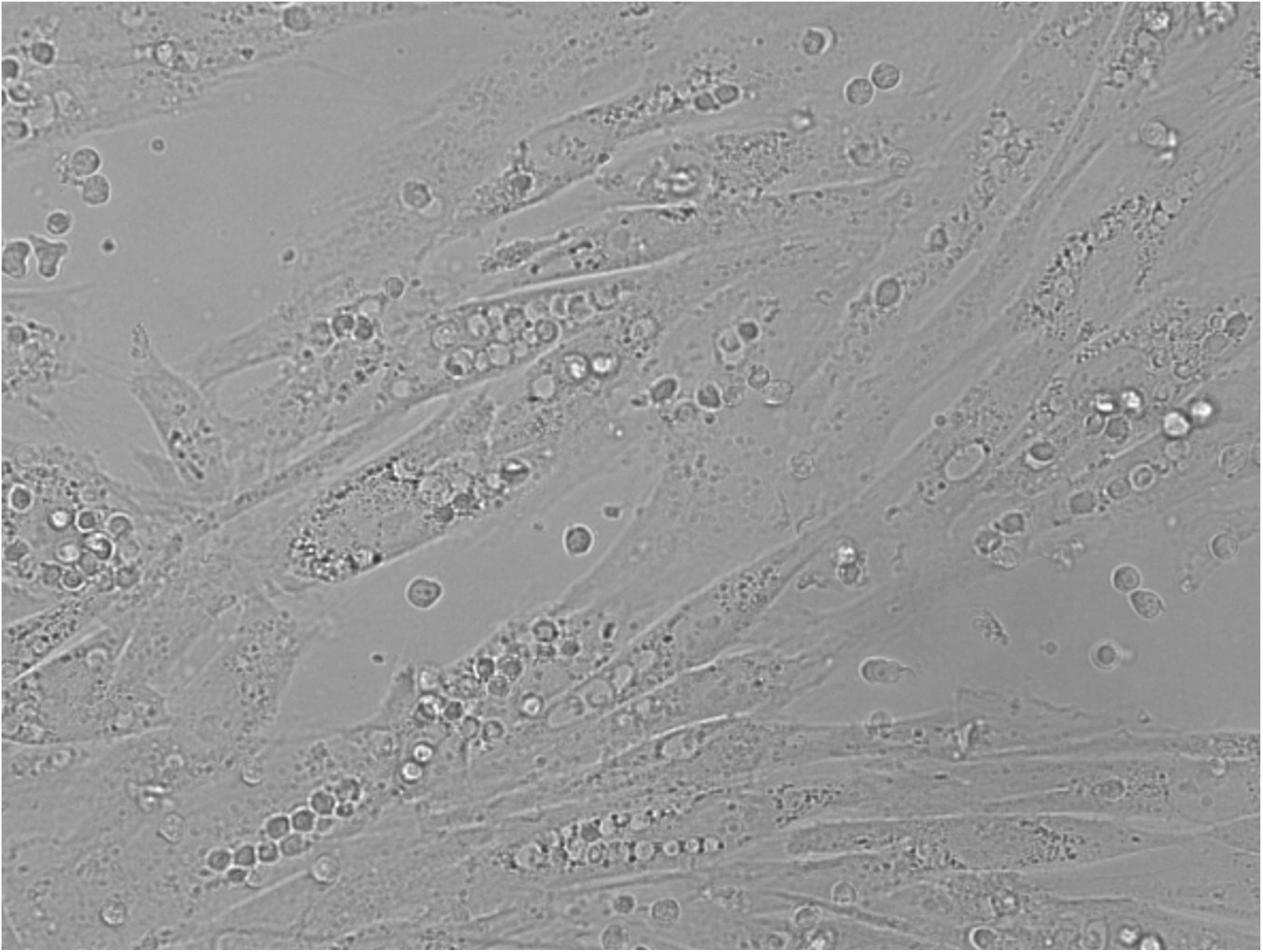
Representative brightfield image of co-culture of NKTert and CLL cells treated with venetoclax.

**Fig. S2:**
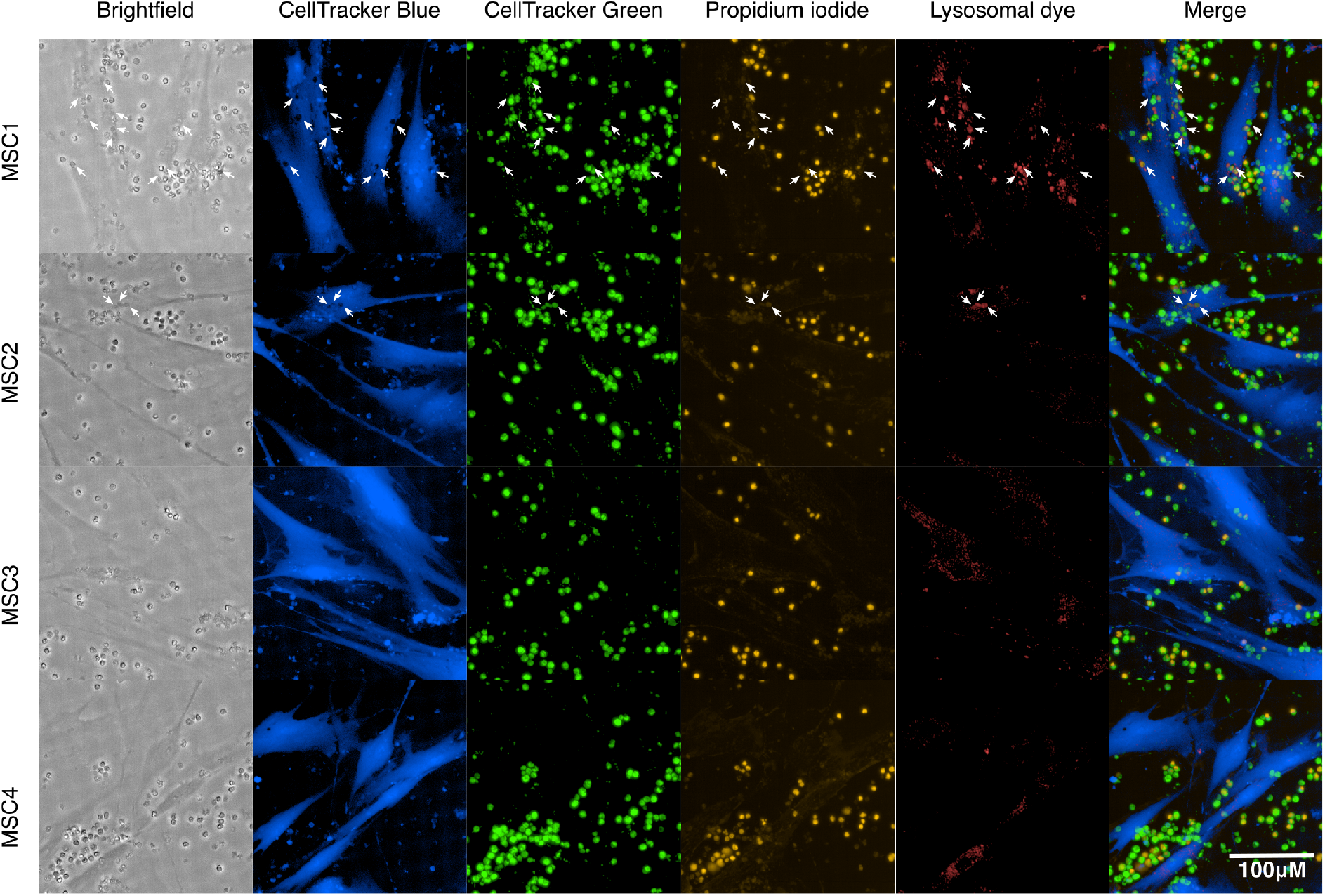
Representative images of mesenchymal stromal cells from four donors (MSC1-4) in co-culture with CLL cells, treated with venetoclax. White arrows indicate all detected phagosomes.

## Movies

**Movie M1: NKTert cell containing phagocytosed CLL cells.** 3D view of confocal microscopy (63x objective) of co-culture of CellTracker Blue labelled NKTert and CellTracker Green labelled CLL cells treated with venetoclax. Green = CellTracker Green, blue = CellTracker Blue, yellow = propidium iodide, red = lysosomal dye NIR, black and white = bright field image.

**Movie M2: MSC cell containing phagocytosed CLL cells.** 3D view of confocal microscopy (63x objective) of co-culture of CellTracker Blue labelled MSC cell from donor MSC1 and CellTracker Green labelled CLL cells treated with venetoclax. Green = CellTracker Green, blue = CellTracker Blue, yellow = propidium iodide, red = lysosomal dye NIR.

## Notes

### Competing Interest Statement

The authors have declared no competing interest.

## References

Baccin C, Al-Sabah J, Velten L, Helbling PM, Grünschläger F, Hernández-Malmierca P, Nombela-Arrieta C, Steinmetz LM, Trumpp A, Haas S. 2020. Combined single-cell and spatial transcriptomics reveal the molecular, cellular and spatial bone marrow niche organization. Nat Cell Biol 22:38–48.

Balakrishnan K, Peluso M, Fu M, Rosin NY, Burger JA, Wierda WG, Keating MJ, Faia K, O’Brien S, Kutok JL, Gandhi V. 2015. The phosphoinositide-3-kinase (PI3K)-delta and gamma inhibitor, IPI-145 (Duvelisib), overcomes signals from the PI3K/AKT/S6 pathway and promotes apoptosis in CLL. Leukemia 29:1811–1822.

Baryawno N, Przybylski D, Kowalczyk MS, Kfoury Y, Severe N, Gustafsson K, Kokkaliaris KD, Mercier F, Tabaka M, Hofree M, Dionne D, Papazian A, Lee D, Ashenberg O, Subramanian A, Vaishnav ED, Rozenblatt-Rosen O, Regev A, Scadden DT. 2019. A Cellular Taxonomy of the Bone Marrow Stroma in Homeostasis and Leukemia. Cell 177:1915–1932.e16.

Cheng S, Ma J, Guo A, Lu P, Leonard JP, Coleman M, Liu M, Buggy JJ, Furman RR, Wang YL. 2014. BTK inhibition targets in vivo CLL proliferation through its effects on B-cell receptor signaling activity. Leukemia 28:649–657.

Choi MY, Kashyap MK, Kumar D. 2016. The chronic lymphocytic leukemia microenvironment: Beyond the B-cell receptor. Best Pract Res Clin Haematol 29:40–53.

Ding L, Zhang W, Yang L, Pelicano H, Zhou K, Yin R, Huang R, Zeng J. 2018. Targeting the autophagy in bone marrow stromal cells overcomes resistance to vorinostat in chronic lymphocytic leukemia. Onco Targets Ther 11:5151–5170.

Dogusan Z, Montecino-Rodriguez E, Dorshkind K. 2004. Macrophages and stromal cells phagocytose apoptotic bone marrow-derived B lineage cells. J Immunol 172:4717–4723.

Fiorcari S, Brown WS, McIntyre BW, Estrov Z, Maffei R, O’Brien S, Sivina M, Hoellenriegel J, Wierda WG, Keating MJ, Ding W, Kay NE, Lannutti BJ, Marasca R, Burger JA. 2013. The PI3-kinase delta inhibitor idelalisib (GS-1101) targets integrin-mediated adhesion of chronic lymphocytic leukemia (CLL) cell to endothelial and marrow stromal cells. PLoS One 8:e83830.

Kanehisa M, Goto S. 2000. KEGG: Kyoto Encyclopedia of Genes and Genomes. Nucleic Acids Res 28:27–30.

Kawano Y, Kobune M, Yamaguchi M, Nakamura K, Ito Y, Sasaki K, Takahashi S, Nakamura T, Chiba H, Sato T, Matsunaga T, Azuma H, Ikebuchi K, Ikeda H, Kato J, Niitsu Y, Hamada H. 2003. Ex vivo expansion of human umbilical cord hematopoietic progenitor cells using a coculture system with human telomerase catalytic subunit (hTERT)-transfected human stromal cells. Blood 101:532–540.

Kurtova AV, Balakrishnan K, Chen R, Ding W, Schnabl S, Quiroga MP, Sivina M, Wierda WG, Estrov Z, Keating MJ, Shehata M, Jäger U, Gandhi V, Kay NE, Plunkett W, Burger JA. 2009. Diverse marrow stromal cells protect CLL cells from spontaneous and drug-induced apoptosis: development of a reliable and reproducible system to assess stromal cell adhesion-mediated drug resistance. Blood 114:4441–4450.

Lagneaux L, Delforge A, Bron D, De Bruyn C, Stryckmans P. 1998. Chronic lymphocytic leukemic B cells but not normal B cells are rescued from apoptosis by contact with normal bone marrow stromal cells. Blood 91:2387–2396.

Ritchie ME, Phipson B, Wu D, Hu Y, Law CW, Shi W, Smyth GK. 2015. limma powers differential expression analyses for RNA-sequencing and microarray studies. Nucleic Acids Res 43:e47.

Subramanian A, Tamayo P, Mootha VK, Mukherjee S, Ebert BL, Gillette MA, Paulovich A, Pomeroy SL, Golub TR, Lander ES, Mesirov JP. 2005. Gene set enrichment analysis: a knowledge-based approach for interpreting genome-wide expression profiles. Proc Natl Acad Sci U S A 102:15545–15550.

Ten Hacken E, Burger JA. 2016. Microenvironment interactions and B-cell receptor signaling in Chronic Lymphocytic Leukemia: Implications for disease pathogenesis and treatment. Biochim Biophys Acta 1863:401–413.

Zhang W, Trachootham D, Liu J, Chen G, Pelicano H, Garcia-Prieto C, Lu W, Burger JA, Croce CM, Plunkett W, Keating MJ, Huang P. 2012. Stromal control of cystine metabolism promotes cancer cell survival in chronic lymphocytic leukaemia. Nat Cell Biol 14:276–286.

